# Response of human iPSC-cardiomyocytes to adrenergic drugs assessed by high-throughput pericellular oxygen measurements and computational modeling

**DOI:** 10.1101/2025.06.27.662066

**Authors:** Weizhen Li, M. Amin Forouzandehmehr, David McLeod, Maria R. Pozo, Yuli W. Heinson, Matthew W. Kay, Zhenyu Li, Stefano Morotti, Emilia Entcheva

## Abstract

Rate- and contractility-modulating drugs, such as adrenergic agonists and antagonists, are widely used in the treatment of cardiovascular conditions. Preclinical assessment of new modulators of rate, inotropy and metabolism can be aided by high-throughput (HT) methods for chronic measurements, coupled with scalable human induced pluripotent stem cell-derived cardiomyocytes (hiPSC-CMs). Here, we evaluate the utility of long-term optical (label-free) measurements of pericellular oxygen in a HT format (96-well plates) for the assessment of the effectiveness of adrenergic drugs in hiPSC-CMs. Quantitative oxygen consumption metrics were derived and correlated to measurements performed in the same samples using all-optical electrophysiology. Adrenergic agonists significantly increased oxygen consumption rate (OCR), best seen in the kinetics of initial depletion of pericellular oxygen, i.e. time to reach 5%. Adrenergic antagonists decreased OCR, best quantified using steady-state values for pericellular oxygen after at least 5 hours. OCR-based drug type identification correlated well with the acute spontaneous rate measurements in the same samples. Direct rate modulation with chronic optogenetic pacing sped up OCR in hiPSC-CMs. Blebbistatin, an excitation-contraction uncoupler, significantly reduced OCR. Computational modeling helped interpret our results by capturing the effects of pacing rate, adrenergic stimulation, and blebbistatin on oxygen consumption, thereby highlighting the key contribution of inotropy and mechanical contraction to OCR in hiPSC-CMs. We conclude that HT label-free optical oxygen measurements and the comprehensive *in silico* hiPSC-CM models, constrained by such measurements, represent valuable human-based approaches for non-invasive assessment of rate- and metabolism-modulating drugs in preclinical studies.

## INTRODUCTION

Modulation of heart rhythm by the sympathetic nervous system (SNS) through action on the adrenergic receptors is an essential physiological response. Catecholamines like norepinephrine and epinephrine target adrenergic receptors to trigger systemic fight-or-flight response, which includes increase in heart rate, contractility and energy consumption. In 1948, Ahlquist [1] categorized the adrenergic receptors into two distinct types, α- and β-adrenergic, based on their pharmacological responsiveness and sensitivity to stimulation by various catecholamines. Subsequent studies have identified them as G-protein coupled receptors (GPCRs)[2, 3], and divided them into nine subtypes (α1A, α1B, α1D, α2A, α2B, α2C, β1, β2, β3)[4]. The adrenoreceptor subtypes most prevalent in the human heart are β_1_ and β_2,_ with lower presence of α1 subtypes [4–6]. The physiological activation of cardiac adrenoreceptor α_1_ and β_1_ yields immediate positive inotropic and chronotropic effects (β_1 >>_ α_1_), increasing cardiac output, as well as leading to cardiac hypertrophy over prolonged action. Conversely, adrenergic antagonists slow down heart rate and overall metabolism.

Clinically, the regulation of cardiac adrenoreceptors has broad applications and therefore the development of respective pharmacological agents has been of great interest. Beta-adrenergic antagonists/blockers are among the most successful cardiac drugs in use[7]. Currently, there are over a dozen β-adrenergic blockers approved by the FDA for various cardiac conditions, including hypertension, angina pectoris, cardiac arrhythmias, hypertrophic cardiomyopathy, and heart failure, among others. Beta-blockers are also used to treat non-cardiac conditions, such as glaucoma and muscle tremors. On the other hand, β-adrenergic agonists act similarly to norepinephrine and epinephrine, increasing cardiac contractility, muscle performance and heart rate. They are clinically used to treat bradycardia, asthma, chronic obstructive pulmonary disease, and allergic reactions[8]. There is interest in developing receptor specific rate-modulating drugs. Therefore, high-throughput methods for reliable and fast preclinical testing of candidate compounds and quantitative methods for comprehensive mechanistic assessment are desirable.

Human-induced pluripotent stem cell-derived cardiomyocytes (hiPSC-CMs) offer scalability and relevance to human physiology. These characteristics make them an attractive model for high-throughput (HT) preclinical drug cardiotoxicity screening [9]. In addition, hiPSC-CMs derived from patients with genetic cardiac diseases can recapitulate aspects of patient’s specific response to pharmacological perturbations [10]. This scalable experimental model of the human heart can be coupled with HT technology for drug screening applications, offering new options for the human-centered modernization proposed by the FDA and NIH [11]. Electromechanical activity in cardiomyocytes is the major contributor to their oxygen consumption [12]. This study aimed to assess the applicability of longitudinal HT optical imaging of pericellular oxygen (in 96-well plates)[13, 14] as a label-free method to detect the action of adrenergic drugs in human iPSC-CMs, cross-validated with measurements of action potentials and spontaneous rates using all-optical electrophysiology [15] in the same samples. To further advance the mechanistic understanding of how adrenergic drugs modulate the coupled electrical, mechanical, and metabolic states of human iPSC-CMs, we complemented our experimental studies with computational modeling. This enabled simulation of the multifaceted effects of adrenergic agonists and of excitation-contraction uncouplers on hiPSC-CM electrophysiology, calcium handling, contractility and oxygen consumption (key readouts accessible through our HT all-optical platforms) and facilitated the interpretation of experimental results. Our hybrid experimental and computational new approach methodology (NAM) [11, 16, 17], integrating scalable human iPSC-CMs, HT all-optical experimental platforms, and validated *in silico* tools, provides a powerful framework for accelerating drug development.

## MATERIALS AND METHODS

### Experimental Methods

#### Human iPSC-CM culture

Human iPSC cardiomyocytes (iCell Cardiomyocytes^2^ from CDI/Fujifilm) were used in this study. Cells were thawed as per manufacturer’s protocol and plated at 50,000 cells per 96-well (1.56 x 10^5^ cells / cm^2^) to form a syncytium. We attached oxygen sensors, covering half of each well, to glass-bottom 96-well plates. The sensors were sterilized and fibronectin coated before cell plating, as described previously [13, 14]. Cells were cultured for 5 days with culture medium exchange every other day before experiments. Pre-treatment oxygen consumption was measured a day in advance, after cell medium exchange.

#### Treatment with adrenergic drugs and excitation-contraction uncouplers

Adrenergic agonists isoproterenol (Sigma-Aldrich I6504) and phenylephrine (Sigma-Aldrich P6126) were used at a dosage of 10 µM applied for 24 hours. Adrenergic antagonists sotalol (Sigma-Aldrich cardiotoxicity ligand set) and propranolol (Sigma-Aldrich P0884) were used at a dosage of 30 µM and 10 µM for 24 hours respectively. Isoproterenol and phenylephrine were diluted in sterile water (Sigma-Aldrich W4502). Sotalol and propranolol were prepared in DMSO (Invitrogen D12345) with a final concentration of 0.3% and 0.1%. As a negative control, we incorporated samples treated with 0.3% DMSO for comparison. All prepared drug solutions were filtered with 0.2 µm sterile filter (VWR 101093-372) before application. Concentrations for the adrenergic drugs were chosen based on prior studies summarized in **Table 1**. An excitation-contraction uncoupler (blebbistatin, BLEBB) was applied at 2.5μM for a subset of experiments. Culture medium volume was 200 µL for all samples.

**Table 1.** Adrenergic agonists and antagonists used in human iPSC-CMs.

|  | drugs | target receptors | dose used | reported dosages | notes | ref |
| --- | --- | --- | --- | --- | --- | --- |
| agonists | (-)-isoproterenol hydrochloride | $\beta$ | 1-10 $\mu\text{M}$ | 1 $\mu\text{M}$ | increased calcium transient amplitude; increased max rate of rise and decay | [53] |
| | | | | 0.1-10 $\mu\text{M}$ | increased beat rate in 30min, recovered at 24h | [37] |
| | | | | 1 $\mu\text{M}$ | positive chronotropic effect (increased beat rate) | [54] |
| | | | | 50 $\mu\text{M}$ | increased conduction velocity & increased contraction rate in 15min | [55] |
| | (R)-(-)-phenylephrine hydrochloride | $\alpha_1$ | 10 $\mu\text{M}$ | 1 $\mu\text{M}$ 48h | iPSC-CM maturation (increase in cell area, organized sarcomeres, hypertrophy marker ANF) | [56] |
| | | | | 1 $\mu\text{M}$ 48h | hypertrophic cardiomyopathy conditioning | [57] |
| | | | | 100 $\mu\text{M}$ 1h | induced Takotsubo syndrome (TTS)-like heart dysfunctions (APD-prolongation and Vmax-suppression) | [58] |
| antagonists | sotalol | $\beta$ | 30 $\mu\text{M}$ | 30-500 $\mu\text{M}$ | decreased beat rate at 30 $\mu\text{M}$; FPDc prolongation at 100 to 500 $\mu\text{M}$ | [59] |
| | | | | 0.1-100 $\mu\text{M}$ | decreased beat rate; APD prolongation | [60] |
| | | | | 10-300 $\mu\text{M}$ | inter-individual differences in response to sotalol | [61] |
| | (±)-propranolol hydrochloride | $\beta$ | 10 $\mu\text{M}$ | 10 $\mu\text{M}$ | inhibition of $I_{\text{NaL}}$ | [62] |
| | | | | 1-10 $\mu\text{M}$ | decreased beat rate, decreased sarcomere shortening; minimal effect on duration of contraction and relaxation, except at high doses. | [63] |
| | | | | 3 $\mu\text{M}$ | decreased beat rate | [64] |
|  |  |  |  | NA | decrease in beat rate of hiPSC-CM spheroids; no effect on the total cAMP production | [65] |

#### Optical measurements of pericellular oxygen and data analysis

Pericellular oxygen was measured using Ruthenium-based oxygen sensing film and a camera system (PreSens, Germany) in the 37°C humidified incubator with 5% CO_2_, as described previously [13, 14]. The exposure time was set to 0.8 s and oxygen images were taken every 5 min in this study. The measurement is based on fluorescence quenching by oxygen and absolute values of pericellular oxygen can be extracted from the acquired images based on the Stern-Volmer equation and a two-point calibration procedure [13, 18]. Time-dependent oxygen calibration was applied in the first two hours of recording, considering temperature and humidity changes from ambient to incubator. Whole-plate data normalization was applied to avoid negative values in the oxygen reading and to set the maximum oxygen reading to be equilibrium oxygen concentration in 37°C, 5% CO_2_ humidified incubator, at 18.6% [12].

#### High throughput all-optical electrophysiology

Voltage and calcium transients were measured optically using fluorescent reporters and a high-throughput whole plate imaging [15, 19]. After adrenergic treatment, samples were labeled with membrane voltage dye BeRST1 at 1 μM (Ex/Em 660/680nm, from Evan W. Miller, University of California, Berkeley) and calcium dye Rhod-4 AM at 10 μM (Ex/Em 530/590nm, AAT Bioquest). The system included excitation filters in front of the LEDs (AT655/30m and ET525/30m, Chroma, Bellows Falls, VT) and multi-bandpass emission filter was used (ET595/40m + 700lp, Chroma). Further details on LED specifications, dichroic mirrors and filters for the design of this system can be found in our prior publication. Signals were acquired at 100 frames per second and transients were extracted and analyzed, as done previously [20]. These electromechanical measurements were done 24 h after the initial application of the adrenergic modulators and with re-application immediately before the recordings.

#### In-incubator optical pacing with pericellular oxygen monitoring

To enable optical pacing, hiPSC-CMs were infected with adenovirus Ad-CMV-hChR2-eYFP [21] five days after cell plating, and drug treatment was done two days later. A blue LED (470 nm) light source was mounted above the PreSens oxygen measuring system in the incubator. The illuminated area was adjusted to cover 12 samples at intensity 0.13-0.3 mW/ mm^2^. Optical pacing was conducted for about 5 hours at 1Hz with 10ms pulse duration, and optical sensing of oxygen was continuous with images acquired every 2 min.

#### Immunolabeling and cell imaging

Monoclonal anti-α-actinin primary antibody (Sigma-Aldrich A7811) and goat anti-mouse Alexa fluor 647 secondary antibody (ThermoFisher A21235) were used for human iPSC-CMs alpha actinin labeling. Hoechst 33342 (ThermoFisher 62249) was used for labeling the cell nuclei. Human iPSC-CMs on glass bottom 96-well plate were imaged with an inverted Zeiss LSM 710 microscope. Human iPSC-CMs grown on oxygen sensors were imaged with Leica TCS SP8 microscope with a 25x water immersion objective.

#### Statistical analysis

Statistical analyses were performed with GraphPad Prism 7. Grouped bar-dot plots were presented as mean ± SD with individual data points. Ordinary one-way ANOVA with Dunnett’s correction for group-wise comparison were applied to identify statistical significance between the control group and treatment groups. Linear regression was applied when correlating the pericellular oxygen results and the electrophysiological measurements.

### Computational Methods

#### Improved estimation of oxygen consumption from hiPSC-CM simulations

We complemented our experimental investigation by performing a computational analysis with a model of electro-mechano-energetic coupling in hiPSC-CMs [22], recently developed based upon the established lineage of *in silico* models by Paci and colleagues [22–24], **Table 2**. Here, we modified the estimation of oxygen consumption rate (OCR) in [22] to: a) account for the SERCA contribution to energy consumption, and b) accurately recapitulate the predominant contribution of myofilament ATPase to the total OCR [25–27]. Note that these changes do not affect the electrical and mechanical behavior of the published hiPSC-CM model. In the updated version, OCR is modeled as the sum of oxygen tied to ATP demand from several processes using the following formulation:

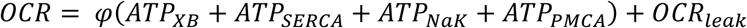

**Table 2.** Computational modeling of hiPSC-CMs with electro-mechano-metabolic responsiveness: evolution of models and summary of key modifications

|  | Reference | Model component | Modifications |
| --- | --- | --- | --- |
| 1 | [66] | Metabolite-sensitive SERCA pump |  |
| 2 | [67] | Metabolite-sensitive myofilament |  |
| 3 | [23] | Human iPSC-CM model (electrical) |  |
| 4 | [24] | Human iPSC-CM model (electrical + mechanical) | Integrating [1], [2], [3] into a new hiPSC-CM model |
| 5 | [22] | Human iPSC-CM model (electrical + mechanical + metabolic) | Introducing oxygen consumption rate (OCR) into [4] |
| 6 | ***This study*** | Human iPSC-CM model with improved OCR, modeling of adrenergic agents and excitation-contraction uncouplers | * Improving [5] to better match experimental OCR data in hiPSC-CMs<br>* Empirically-constrained modeling of ISO in hiPSC-CMs<br>* Empirically-constrained modeling of BLEBB in hiPSC-CMs |

Each term represents the oxygen equivalent of the ATP flux consumed by the respective cellular processes. ATP_XB_, ATP_SERCA_, ATP_NaK_, ATP_PMCA_ are the ATP consumption rates for myofilament, SERCA pump, sodium-potassium ATPase, and plasmalemmal calcium pump (PMCA), respectively. OCR_leak_ represents non-phosphorylating respiration (proton leak + maintenance). The stoichiometric conversion factor 𝜑 converts ATP hydrolysis to oxygen, (i.e., oxygen per ATP) and was considered equal to 0.2 [28, 29]. ATP utilization rates (in mM/s) were computed as follows:

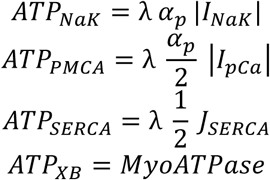

I_NaK_ represents the sodium-potassium ATPase current, I_pCa_ denotes the PMCA current, J_SERCA_ is the sarcoplasmic reticulum (SR) calcium uptake flux, and *MyoATPase* refers to the myosin ATPase activity of cross-bridges (XB) in the contractile apparatus. The scaling factor 𝛼_𝑝_ (equal to 1.1626 × 10^−1^ mM·F·s⁻¹·A⁻¹) converts current density to concentration change rates [23] and is calculated as 𝛼_𝑝_ = Cm/(F·Vc·10⁻¹⁸), using cell volume (Vc) and Faraday’s constant (F), and membrane capacitance (Cm). The scaling factor λ (equal to 0.008) was applied to non-contractile OCR pathways to achieve the physiologically observed ∼70% contribution of myofilament ATP_XB_ to total OCR during spontaneous beating [25, 26]. OCR_leak_ was set to 0.01 mM/s to obtain a contribution of non-phosphorylating respiration to total OCR between 10 and 25%, consistent with experimental ranges determined in hiPSC-CMs [30, 31].

The updated model, simulated using MATLAB R2025a (MathWorks, Natick, MA, USA), was used to analyze hiPSC-CMs’ electrophysiology, mechanics, and energetics at steady-state during spontaneous activity or during pacing at variable rates. We also simulated the experimental protocol characterizing dynamics of pericellular oxygen depletion over 5 hours (during spontaneous activity). Here, the initial pericellular oxygen concentration was set to 0.1772 mM to match the corresponding value of 18.5% that is the maximum concentration obtained in the incubator. Variation in pericellular oxygen concentration over time is calculated as a first-order process following Fick’s law:

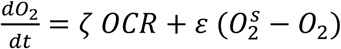

Where *O*_2_^s^ represents the source (bath) oxygen concentration and is assumed to be constant (equal to 0.1772 mM). The pericellular scaling factor 𝜁 accounted for the ratio of total cell volume to well volume was equal to 0.002. The scaling factor ε was set to 2.56 × 10^−4^ s^-1^ to reproduce the 5-hour decay observed in control experiments in this study.

#### Computational modeling of hiPSC-CMs’ response to adrenergic agents (isoproterenol)

We simulated the steady-state effects of 10 𝜇M isoproterenol (ISO) administration by modifying key model parameters corresponding to functional changes induced by protein kinase A (PKA) phosphorylation on multiple cellular targets (**Table 3A**). In line with previous modeling studies [32–34], each change was selected to reproduce the responses measured in hiPSC-derived and (wherever not available) human ventricular cardiomyocytes. The overall effect of adrenergic stimulation was validated against ISO-induced changes in emerging cellular properties such as beat rate, action potential and calcium transient duration experimentally measured in this study and others [35–37], as reported in **Table 3B**.

**Table 3.** Computational modeling of hiPSC-CMs and the effects of isoproterenol (ISO): A. summary of changes in model parameters to capture PKA phosphorylation and ISO effects compared to control (CON); B. comparison of biomarkers between model generated data and experimental *in vitro* data for hiPSC-CMs, including this study.

| **A** | **PKA Target** | **Reference** | **PKA phosphorylation effect** | **Change in ISO vs. CON** |
| --- | --- | --- | --- | --- |
| | L-type $\text{Ca}^{2+}$ current ($I_{\text{CaL}}$) | [68] | - Increased maximal conductance | GCaL: +30% |
| | Slow delayed rectifier $\text{K}^{+}$ current ($I_{\text{Ks}}$) | [69] | - Increased maximal conductance | GKs: +50% |
| | Rapid delayed rectifier $\text{K}^{+}$ current ($I_{\text{Kr}}$) | [33] | - Increased maximal conductance | GKr: +30% |
| | Ryanodine receptor, SR $\text{Ca}^{2+}$ release flux ($J_{\text{rel}}$) | [70] | - Faster opening: reducing Po half activation constant and,<br>- increased maximal release gain. | RyRohalf: -30%<br>$J_{\text{rel}}\text{max}$: +30% |
| | Funny current ($I_{\text{f}}$) | [71] | - Depolarized shift of half-activation voltage.<br>- Increased maximal conductance | $V_{1/2}$: +8 mV<br>GI<sub>f</sub>: +10% |
| | $\text{Na}^{+}/\text{K}^{+}$ ATPase current ($I_{\text{NaK}}$) | [72] | - Increased $I_{\text{NaK}}$ affinity for intracellular $\text{Na}^{+}$ | Km_Na: -10% |
| | SERCA pump, SR $\text{Ca}^{2+}$ uptake ($J_{\text{up}}$) | [73] | - Increased affinity for intracellular $\text{Ca}^{2+}$<br>- Increased SERCA uptake gain | Km: -30%<br>Vmax: +30% |
| | Myofilament troponin I | [45] | - Decreased sensitivity to intracellular $\text{Ca}^{2+}$ | Kon: -10%<br>KoffL: +10%<br>KoffH: +10%<br>Perm50: +10% |
|  | Myosin-binding protein-C (MyBP-C) | [45]<br>[74] | - Faster cross-bridge recruitment and turnover | fapp: +30% |
|  | Titin phosphorylation | [75] | - Reduced titin passive stiffness → lower passive tension | PconTitin: -10%<br>PexpTitin: -10% |

| **B** | **Biomarker (ISO vs. CON)** | ***In silico*** | ***In vitro* iPSC-CM** | **Source** |
| --- | --- | --- | --- | --- |
| 1 | Increase in beat rate (%) | 24 ▲ | 20 median/ 59 mean ▲<br>42 ▲<br>28 ▲ | 10 $\mu\text{M}$, ***This study***<br>10 $\mu\text{M}$, [37]<br>2 $\mu\text{M}$, [36] |
| 2 | Increase in peak tension (%) | 52 ▲ | 40 ▲ | 10 $\mu\text{M}$, [35] |
| 3 | Change in APD80 (%) | 8.3 ▲ | NS 9.4 median / 10 mean ▲ | 10 $\mu\text{M}$, ***This study*** |
| 4 | Change in CTD80 (%) | 15.8 ▼ | NS 8.3 median / 7.7 mean ▼ | 10 $\mu\text{M}$, ***This study*** |
| 5 | Change in pericellular $\text{O}_2$ at 5h (%) | 98.6 ▼ | 74 median / 67 mean ▼ | 10 $\mu\text{M}$, ***This study*** |

#### Computational modeling of hiPSC-CMs’ response to excitation-contraction uncouplers (blebbistatin)

We simulated the effect of BLEBB on hiPSC-CMs by implementing the approach described in [38]. **Table 4A** reports the parameter changes associated to BLEBB’s primary mechanism: inhibition of the transition from the pre-power-stroke, weakly bound state (XBpreR) to the strongly bound, force-generating state (XBpostR) by blocking Pᵢ release [39–41]. This slows cross-bridge cycling and reduces the fraction of force-producing heads. The overall effect of BLEBB was validated against experimental data describing changes in myofilament properties and OCR (**Table 4B**).

**Table 4.** Computational modeling of hiPSC-CMs and the effects of blebbistatin (BLEBB): A. summary of changes in model parameters implemented to capture BLEBB effects (compared to CON) as previously done in [38]; B. comparison of biomarkers between model generated data and experimental *in vitro* data for hiPSC-CMs, including this study.

**A**
| Modeling the effects of blebbistatin on hiPSC-CMs:<br>updates to [38] FhiPSC-CM model to match experimental data |  |  |  |  |
| --- | --- | --- | --- | --- |
| Muscle myofilament process | Myofilament model parameter | Scaling factor | Effect | Ref. |
| DRX-SRX ratio | F1 | 5 | Increases recruitment into the pre-rotated (XBpreR) state, biasing heads toward the pre-power-stroke configuration | [39]<br>[40]<br>[41] |
|  | F2 | 0.1 | Reduces detachment from the pre-rotated state, stabilizing weakly bound heads |  |
| XB myosin head isomerization | ap2 | 0.03 | Slows the transition from pre-rotated to force-generating state (blocked P<sub>i</sub> release) |  |
| thin-filament cooperativity of activation | Perm50 | 1.4 | Decrease of Ca<sup>2+</sup> sensitivity and cooperativity of force | [46] |

**B**
|  | Biomarker (BLEBB vs. CON) | Simulation | *In vitro* data |
| --- | --- | --- | --- |
| 1 | Change in pCa (%) | 1.5 ▼ | 2.1 to 5.7 ▼<br>5 μM, [46] |
| 2 | Change in nHill (%) | 10.3 ▼ | 44.8 to 73 ▼<br>5 μM, [46] |
| 3 | Change in peak tension (%) | 79.4 ▼ | ~ 80 ▼<br>hiPSC-CMs, 10 μM [35] |
| 4 | Change in OCR at steady state (%) | 36.3 ▼ | 44 median / 41 mean ▼<br>hiPSC-CMs, 2.5 μM, ***This study*** |

## RESULTS

### Integrated system for HT long-term oxygen measurements with all-optical electrophysiology for acute and chronic assessment of adrenergic agonists and antagonists

Pharmaceutical agents targeting adrenergic receptors are a clinically important drug category in active development, including vasopressors, bronchodilators, and controllers of cardiac rhythm. Considering that about 80% of the cardiomyocyte oxygen consumption is related to mechanical contraction, we tested if measurements of pericellular oxygen can be used to quantify the effectiveness of adrenergic agonists and antagonists. To validate the drug treatments, we used sequential optical measurements of pericellular oxygen followed by all-optical electrophysiology in the same samples (in 96-well glass-bottom plates), as shown in **Fig. 1**. Since the oxygen sensors are non-transparent, we used laser-cut semicircles to cover half of each well and allow unimpeded optical electrophysiology measurements in the other half (**Fig. 2A**).

**Figure 1.**
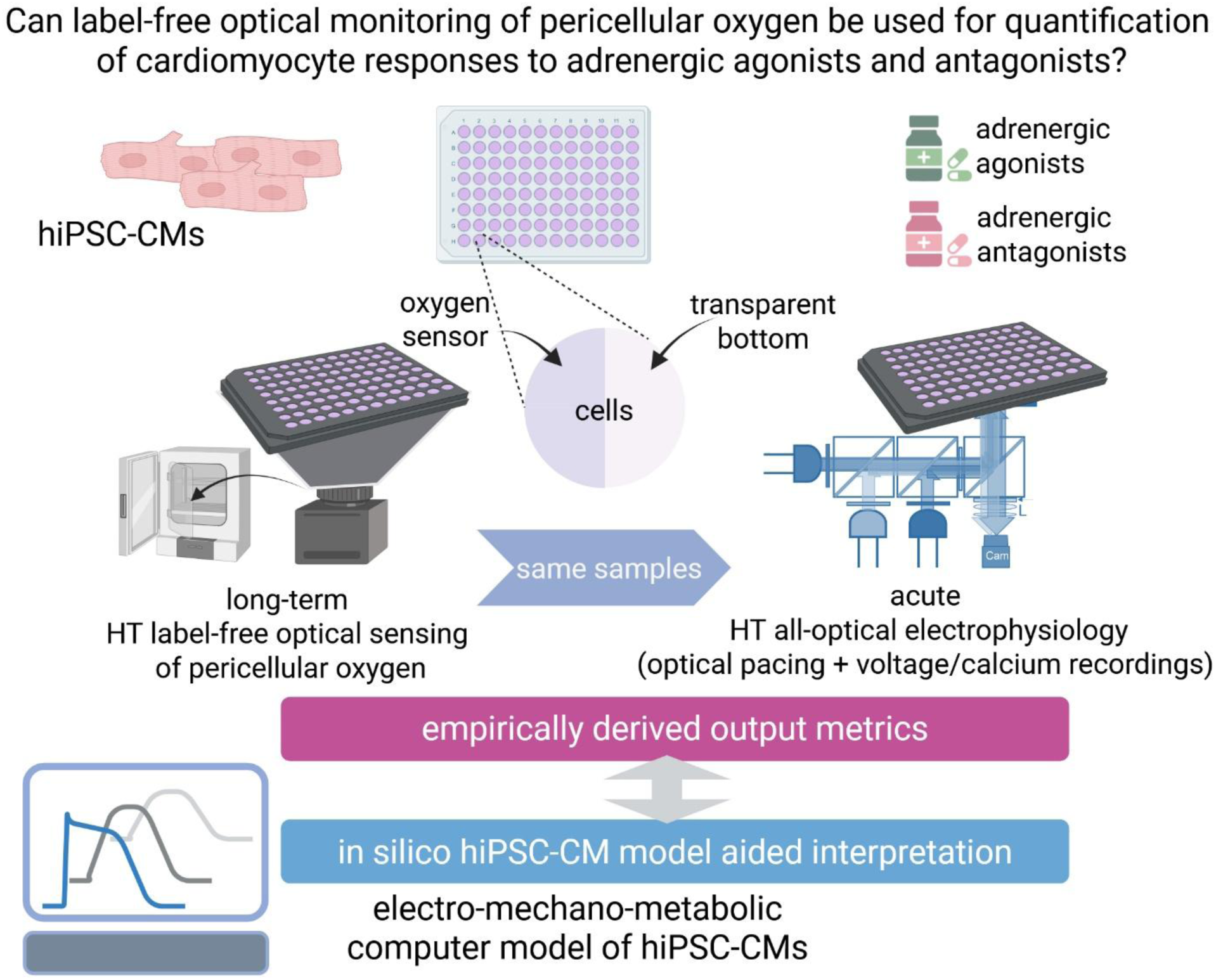
Study overview. Human induced pluripotent stem cell-derived cardiomyocytes (hiPSC-CMs) cultured in 96-well plates can be used for high-throughput (HT) drug testing. This study examines if HT label-free HT optical sensing of peri-cellular oxygen can accurately capture responses of human iPSC-CMs to adrenergic agonists and antagonists and what metrics may be most useful. The study uses acute all-optical electrophysiology in the same samples to register level of activity and correlate to peri-cellular oxygen responses. In addition, computational modeling was used to help investigate the effects of pacing rate, adrenergic agents, and electromechanical uncouplers on oxygen consumption in hiPSC-CMs.

**Figure 2.**
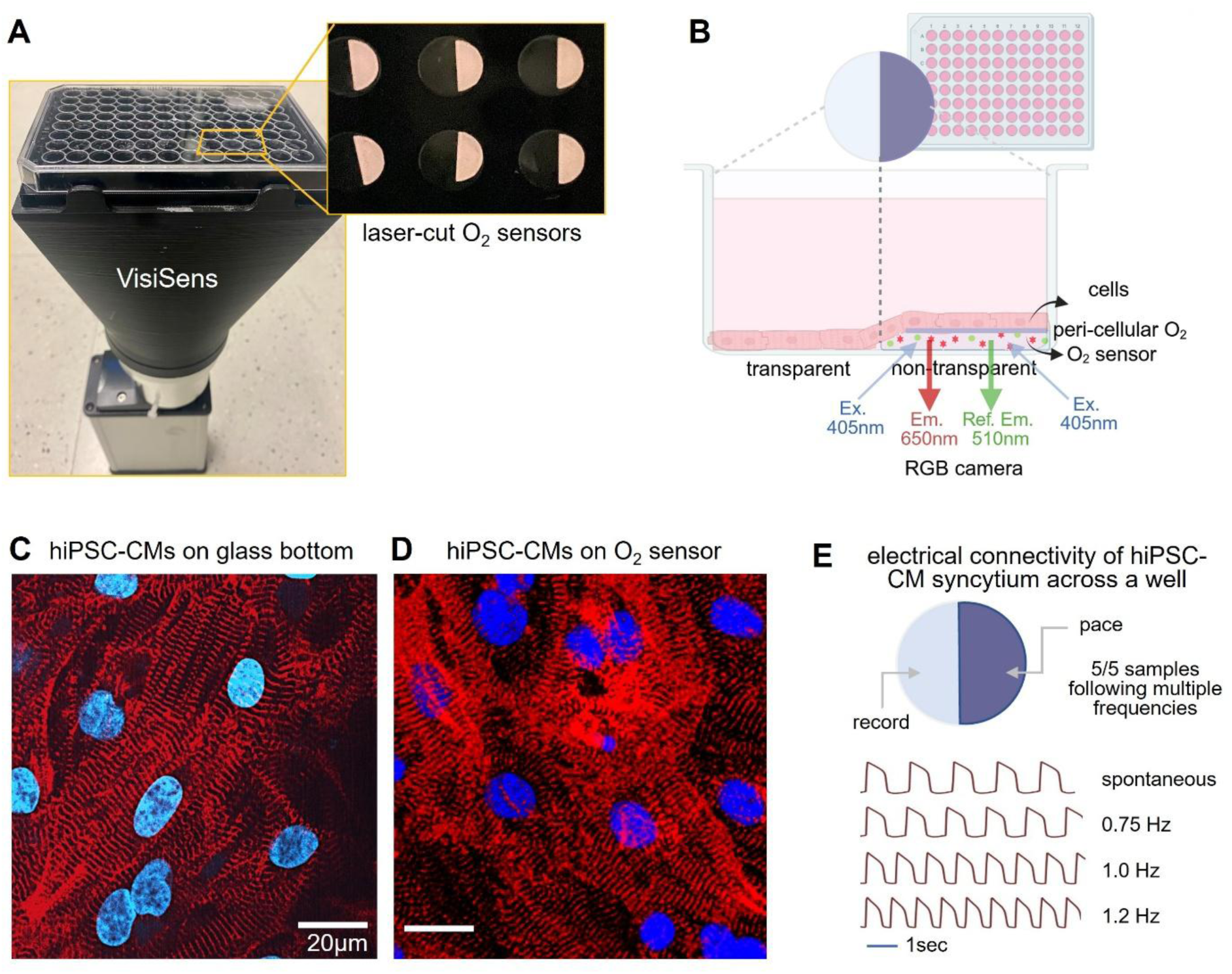
High-throughput optical sensing of pericellular oxygen in standard 96-well plates. **A.** Top: bottom-up view of semi-circular oxygen sensor attached 96 well. Bottom: VisiSens 96-well plate oxygen measuring setup. **B.** Schematic of 96 well peri-cellular oxygen sensing. Cells are grown on the glass bottom on the left side for all-optical electrophysiology measuring, while on the right side, they are grown on the oxygen sensor for real-time oxygen measurements. Oxygen sensitive membrane in the oxygen sensor (pink) contains ruthenium-based oxygen-responsive dye that has fluorescent quenching with peri-cellular oxygen molecules. Under VisiSens 405nm light excitation, the 650nm red fluorescent emission indicates the oxygen level, while a 510nm reference fluorescent emission remains constant regardless of the oxygen level. **C, D.** Human iPSC-CMs cultured on the glass-bottom (C) and oxygen sensor(D). Alpha-actinin signal in red, and nuclei in blue, with 20µm scale bar. **E.** Demonstration of electrical continuity across the two halves of the samples: pacing with different frequencies from an electrode positioned at the sensor side triggers precise responses at the recording site (transparent portion on the left) – shown are example voltage traces (similar results recorded in 5/5 samples).

The oxygen sensor is about 180 µm thick, with an oxygen-sensitive layer, a polyester support layer, and a white optical isolation layer. It has a ruthenium-based oxygen-responsive dye and a reference dye. Cells grow directly on top of the oxygen permeable white optical isolation layer, allowing for continuous pericellular oxygen readings (**Fig. 2B**). Images of hiPSC-CMs grown directly on the glass-bottom of 96-well plate and on the oxygen sensor are shown in **Fig. 2C, D** (alpha actinin in red, nuclei in blue with 20 µm scale bar). Within a well, the densely plated cardiomyocytes formed a contiguous layer (syncytium) across the sensor-covered portion and the other half. Electrical connectivity across the two halves of each well was confirmed by probing with electrical pacing, **Fig. 2E**. Because of the small size, high cell density and electrical connectivity, we assumed homogeneous behavior within a well, off and on the oxygen sensor.

### Adrenergic agonists accelerate oxygen consumption while adrenergic antagonists slow down oxygen consumption; different features of the oxygen depletion curve best differentiate their action from control

Two adrenergic agonists, a β-agonist, isoproterenol (10µM) and an α1-agonist, phenylephrine (10µM), and two β-adrenergic antagonists, sotalol (30µM) and propranolol (10µM), were applied on hiPSC-CMs for up to 24 hours based on the reported studies summarized in **Table 1**. DMSO control with 0.3% DMSO in culture medium was included to consider its potential influence as a solvent. Pre-treatment oxygen consumption showed similar trend across all samples with steep oxygen depletion in the first 3 hours and stabilization after that, as we have shown previously. After 10 hours, some samples exhibited non-monotonic oxygen consumption behavior as we have reported before [13, 14].

Following adrenergic treatment, the initial rate of depletion of pericellular oxygen as well as the steady state levels were greatly affected (**Fig. 3A, B**). **Fig. 3C** summarizes pericellular oxygen levels of hiPSC-CMs at 5 and 22 hours after different treatments. Compared to control, adrenergic agonists accelerated oxygen depletion. The 30 µM sotalol treatment slowed down the initial oxygen consumption (in the first 5 h), but steady state levels (after 22 h) were similar to control, **Fig. 3D**. The 10 µM propranolol treatment slowed down oxygen consumption consistently, evidenced by continuous high pericellular oxygen readings at 5 h and at 22 h. One-way ANOVA with Dunnett’s multiple comparisons suggested statistical significance between control and all treatments (at p<0.01), see figure legend for details.

**Figure 3.**
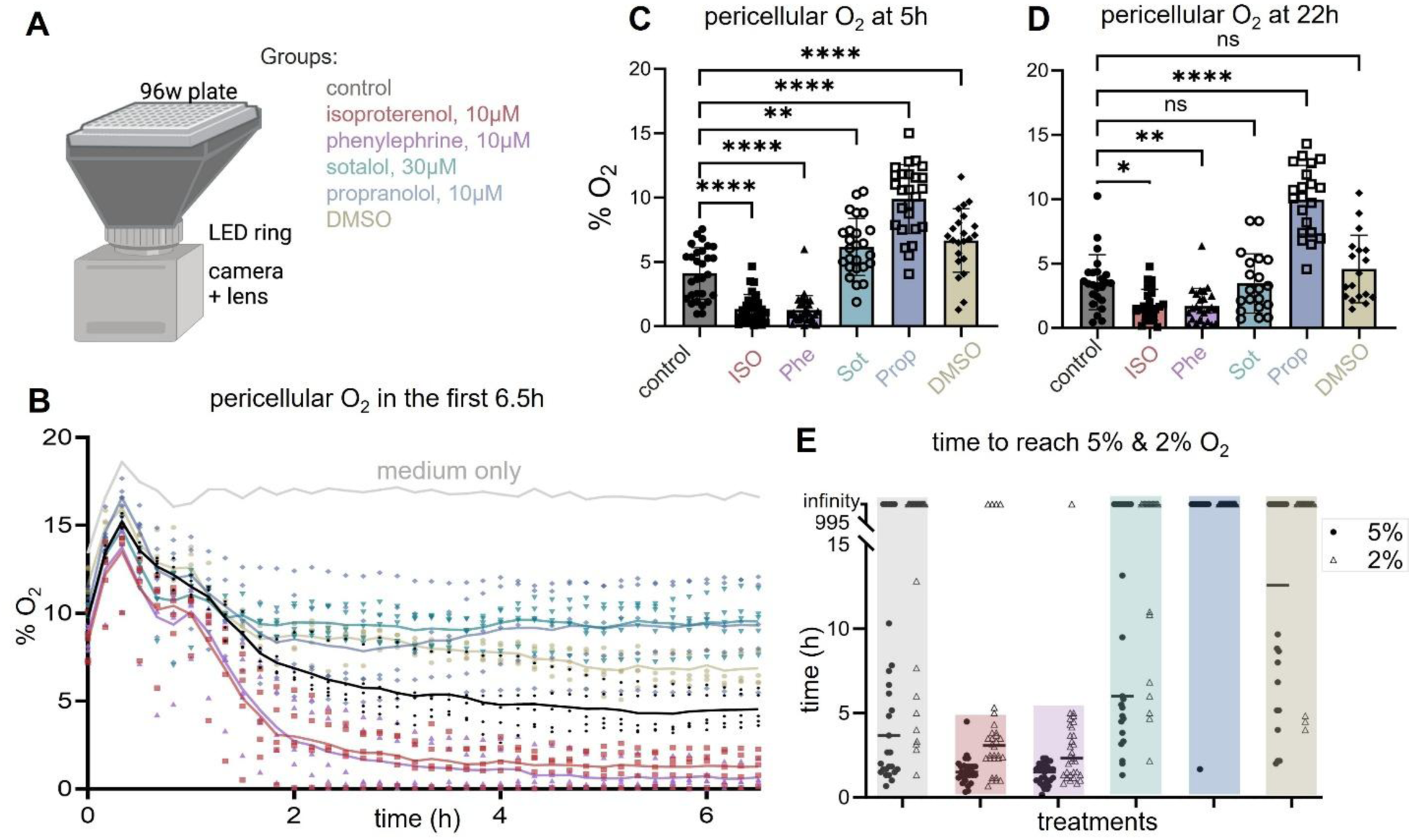
Pericellular oxygen dynamics in response to adrenergic agonists and antagonists. **A.** VisiSens 96-well plate oxygen measurement setup and color code for adrenergic agonists and antagonists. **B.** Pericellular oxygen readout in the first 6.5 hours of treatment with adrenergic agonists and antagonists; data presented as individual samples (wells) along with a trend line representing the average (n=5 per group, with an additional control well with culture medium only, no cells). Drug treatment was done six days after cell plating. **C.** Pericellular oxygen levels at 5 hours post-treatment (n=22-29 per group). One-way ANOVA with Dunnett’s multiple comparisons showed a significant difference between control and sotalol-treated groups (p = 0.001), and between control and all other treatment groups (p < 0.0001). **D.** Pericellular oxygen levels at 22 hours post-treatment (n=18-24 per group). One-way ANOVA with Dunnett’s test revealed significant differences between control and isoproterenol (p = 0.0173), phenylephrine (p = 0.0096), and propranolol (p < 0.0001) treated groups. **E.** Time to reach 5% and 2% pericellular oxygen (n=22-31 per group); some samples never reached below 5%.

To capture the change in rate of depletion we identified two metrics - time to reach 5% and 2% O_2_ - as sensitive to adrenergic modulators (**Fig. 3E**). Adrenergic agonists isoproterenol and phenylephrine accelerated the depletion of pericellular oxygen to below 5% to 1.5 hours in average, and 86% (26 out of 31) of isoproterenol treated and 97% (30 out of 31) of the phenylephrine treated samples reached hypoxic levels - 2% O_2_ – in less than 3 hours after media change. In comparison, the average time to reach 5% O_2_ among the control samples was 4.7 hours and 69% (20 out of 29) didn’t reach 2% O_2_. Among sotalol treated samples 64%(16 out of 25) reached 5% O_2_, and the average time to do so was 6.1 h; 61% of sotalol treated samples (16 out of 26) didn’t reach 2% O_2_. For propranolol treated samples, 7% (2 out of 26) reached 5% O_2_ and none of them reached hypoxic levels, i.e. <2% O_2_. For DMSO control, 54% samples (12 out of 22) reached 5% O_2_ in an average of 6.51 hours, and 83% (19 out of 23) didn’t reach 2% O_2_. Based on previous studies, sotalol (membrane impermeable) is a less potent β-blocker compared to propranolol (membrane permeable), which may explain its milder effects of pericellular oxygen[42].

In summary, adrenergic antagonists (β-blockers) were most easily distinguishable by the higher steady-state values for pericellular oxygen within 5 h post-treatment, while the adrenergic agonists had a distinct signature in the time to reach 5% of pericellular oxygen (significantly faster than other groups).

### Adrenergic perturbations modify electromechanical responses of human iPSC-CMs, including beat rate, action potential and calcium transient morphologies

High-throughput all-optical electrophysiology platform was deployed for electrophysiology functional assessment. Spontaneous activity was documented by measuring membrane voltage and intracellular calcium separately (**Fig. 4A**). Comparison of spontaneous frequency, after normalization by the average rate of control samples, is shown in **Fig. 4B**. On average, 10 μM isoproterenol and 10 μM phenylephrine increased spontaneous rate by 59% and 52% respectively (n=12 to 16 per group). The 30 μM sotalol treatment decreased spontaneous rate by 45% on average (n=12). And the 10 μM propranolol treatment completely suppressed spontaneous activity in 85% (n=16) of the samples. DMSO (at 0.3%) reduced spontaneous activity by 25% on average (n=8). The multiple comparison of one-way ANOVA analysis showed statistically significant difference for isoproterenol, phenylephrine, and propranolol with respect to control (at p<0.02), see figure legend for details.

**Figure 4.**
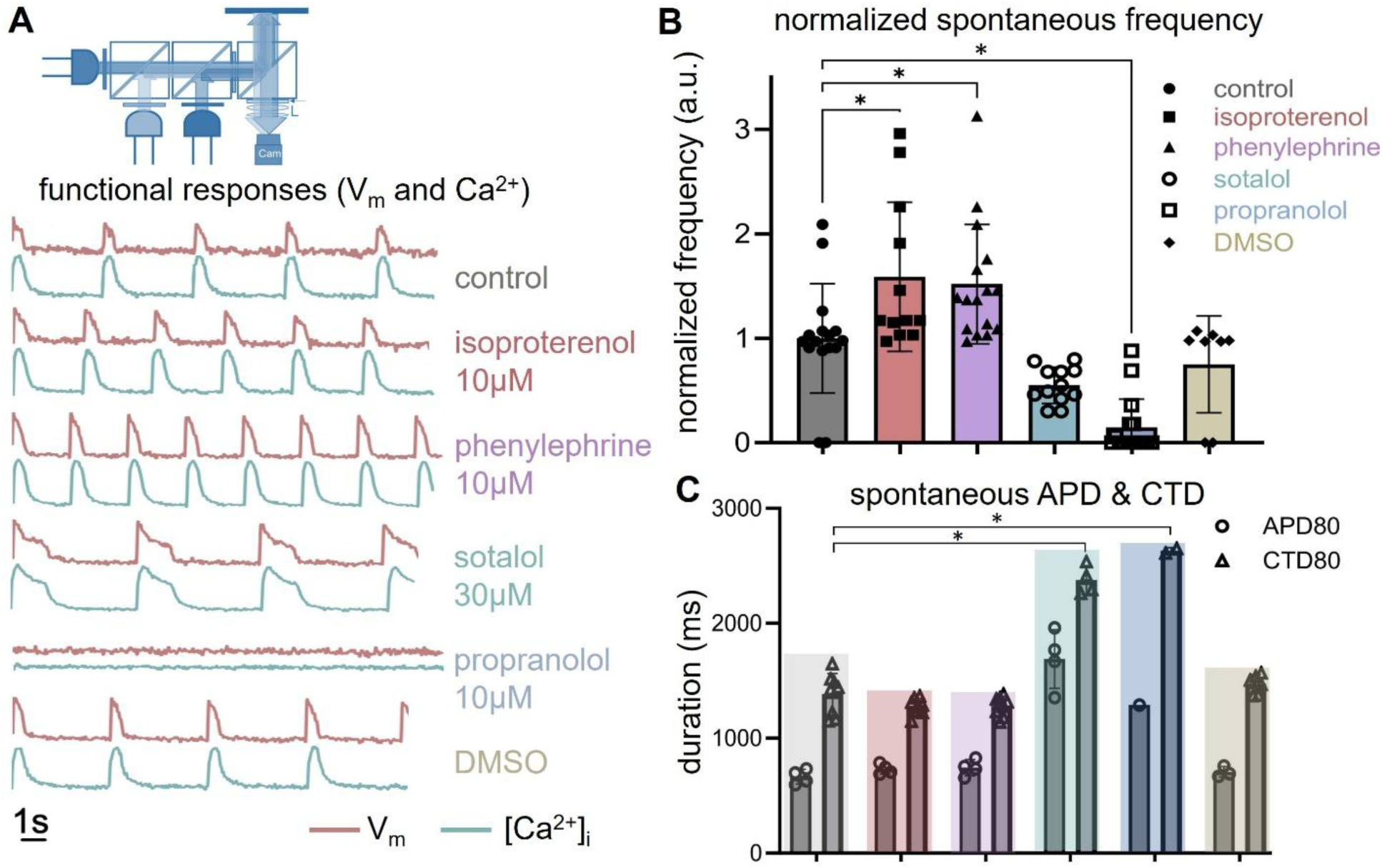
Functional follow-up: voltage and calcium activities in hiPSC-CMs post-adrenergic modulation. **A.** Example traces from the whole-plate all-optical electrophysiology measurement, 24 h after treatment with adrenergic agonists and antagonists. Membrane voltage signal represented in red, while calcium transients are shown in green; these were measured sequentially. **B.** Normalized spontaneous beating frequency of human iPSC-CMs under adrenergic agonist and antagonist treatments (n = 8–16 per group). One-way ANOVA with Dunnett’s multiple comparisons showed significant differences between control and isoproterenol (p = 0.01), phenylephrine (p = 0.0159), and propranolol (p < 0.0001). **C.** Human iPSC-CMs spontaneous action potential duration (APD80) and calcium transient duration (CTD80) under adrenergic agonists and antagonists’ treatments (n=1-8 per group; some samples were completely quiescent). Statistical analysis of CTD80 using ordinary one-way ANOVA and multiple comparisons revealed adjusted p<0.0001 for control vs. sotalol and control vs. propranolol.

We also compared action potential duration (APD80) and calcium transient duration (CTD80), **Fig. 4C**. These rate-uncorrected APD80 and CTD80 showed prolongation in samples treated with adrenergic antagonists compared to the control, with statistically significant difference for sotalol and for propranolol with respect to control (at p<0.001). Sotalol, in particular, is known for APD prolongation caused at least in part by blocking the HERG channel. The adrenergic agonists, as used here, did not significantly affect APD and CTD, except for minor APD80 prolongation combined with CTD80 shortening.

### Correlation between oxygen consumption and electromechanical function in hiPSC-CMs indicates the overall impact of adrenergic perturbations

We sought to uncover potential correlation in oxygen consumption and electromechanical activity in the cardiomyocytes. Pericellular oxygen levels at 5 h after adrenergic treatments were plotted against spontaneous beating frequency within the same samples to probe for such correlation in the iPSC-CMs (**Fig. 5A**). A decrease in 5h pericellular oxygen level correlated with an increase in spontaneous rate. This trend was expected as cardiac electromechanical activity is the dominant process underlying oxygen consumption. Adrenergic antagonist propranolol treated samples had the lowest spontaneous activity level and the highest 5 h pericellular oxygen levels, followed by sotalol treated samples. Propranolol, at the concentration used here, often silenced spontaneous activity and therefore yielded very low oxygen consumption, however, the cells remained viable and excitable. Untreated and DMSO control samples were in the middle of the trend, and adrenergic agonists isoproterenol and phenylephrine presented increased spontaneous activity levels and low 5 h pericellular oxygen levels. Grouping the samples by treatment type, revealed more clearly the trends induced by the adrenergic perturbations (**Fig. 5B**). When this metric was used (pericellular oxygen levels 5 h after treatment), the adrenergic antagonists were clearly identifiable, while the adrenergic agonists, despite the induction of higher rates in the treated samples, were less distinct from controls. The morphology of the response curves can be further used to extract the most sensitive features to correctly identify adrenergic drugs. For example, the time to reach 5% would have been a better metric for adrenergic agonists, as seen in **Fig. 3**.

**Figure 5.**
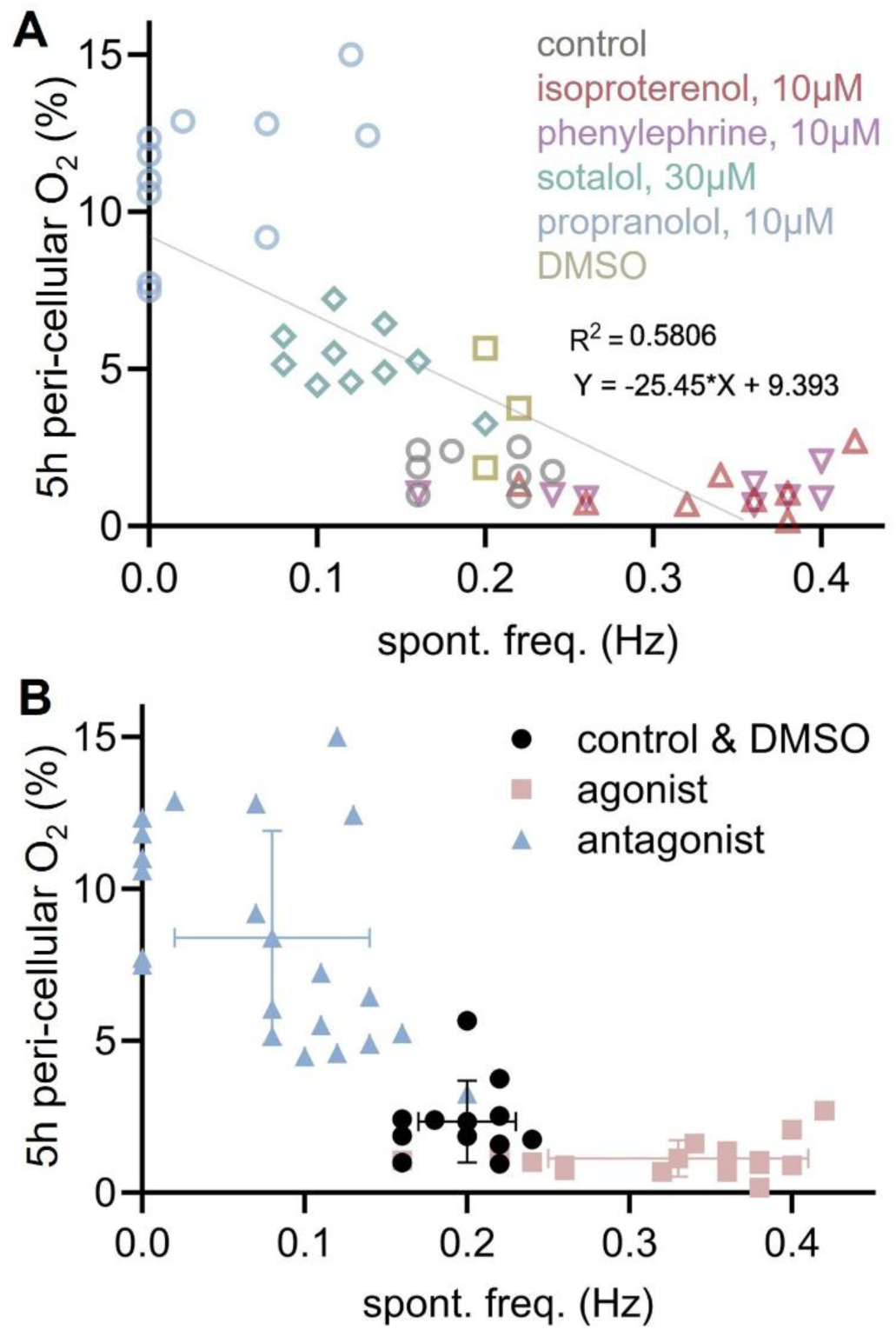
Correlation of peri-cellular oxygen measurements and cardiac cellular activity. **A.** Correlation plot of peri-cellular oxygen readings at 5 h of adrenergic treatment and cell spontaneous frequency (n=3-11 per group, samples colored by treatment). **B.** Grouped correlation plot for adrenergic antagonists, control and adrenergic agonists (n=11-21 per group); bars represent the standard deviations in either parameter per group.

### Active optical pacing directly impacts oxygen consumption. Responses of hiPSC-CMs to adrenergic perturbations are preserved after optogenetic transduction

To confirm the effect of heart rate on oxygen consumption in hiPSC-CMs in a more direct way, we paced the cells directly using optogenetic stimulation. Cells were made light-sensitive through ChR2-eYFP transduction prior to experiments, which we have shown to not cause any electrophysiological side effects[43]. Considering potential interference (by the optogenetic actuator or its reporter) on the optical oxygen measurements, we first confirmed that the modified iPSC-CMs have preserved sensitivity to the adrenergic modulators (**Fig. 6A-B**). Just as seen with unmodified myocytes, adrenergic agonists accelerated initial oxygen depletion (in the first 2 hours), while the application of adrenergic antagonists led to high steady-state levels of pericellular oxygen (around 10%) (n=5 per group).

**Figure 6.**
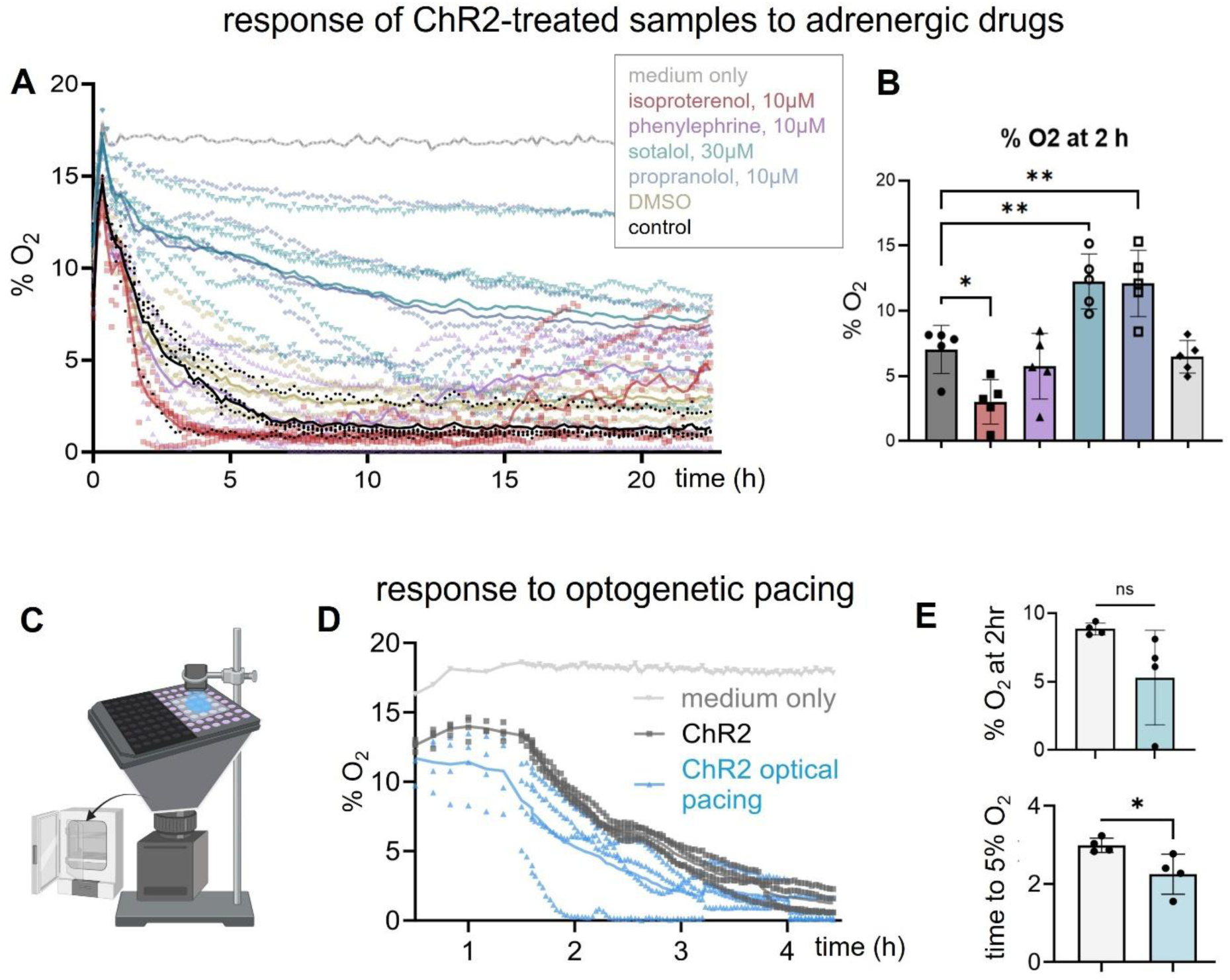
Active perturbation of rate through optogenetic pacing and changes in pericellular oxygen. **A.** Pericellular oxygen of adrenergic treated samples after ChR2 infection (n=5 per group except medium-only group). **B.** Summary of steady-state pericellular oxygen values at 2 h after treatment for all groups. One-way ANOVA with Dunnett’s multiple comparisons showed significant differences between control and isoproterenol (p = 0.0198), sotalol (p = 0.0022), and propranolol (p = 0.0029). **C.** Diagram illustrating the setup for optical pacing within an incubator with peri-cellular oxygen measurement. **D.** Oxygen consumption comparison between optically paced and non-paced samples hiPSC-CMs (n=4 per group). **E.** Comparison of oxygen consumption metrics between optically paced and non-paced samples (top: peri-cellular oxygen level after about 2 h of optical pacing; bottom: time to reach 5% O_2_ level); Unpaired t test was applied for statistical analysis, p = 0.0338.

To test the effects of pacing on oxygen consumption, we deployed the setup, Illustrated in **Fig. 6C**, where a blue LED was used above the 96-well plate for optical pacing, while optical probing of oxygen consumption was done by the system positioned underneath the 96-well plate. Part of the 96-well plate was covered with black tape to block the blue light stimulation and have control (spontaneously beating) samples and actively paced samples. Pacing at 1Hz was chosen as being close to the spontaneous rate of these hiPSC-CMs to not induce severe hypoxia and with high likelihood of capture. In **Fig. 6D**, we show the pericellular oxygen readouts from the optically paced and non-paced samples within 5 h of active optical pacing. Pacing accelerated oxygen consumption and resulted in lower pericellular oxygen readout compared to spontaneously beating control samples in the same plate. The readouts at 2 h are quantified in **Fig. 6E**, where the paced cells have depleted oxygen more than controls, albeit statistical significance was not reached. However, the two groups had statistically different time to reach 5% O_2_ (n=4 per group, p = 0.0338), just as we saw that this metric is better in distinguishing the action of adrenergic agonists (**Fig. 3**). This confirmed similarity between adrenergic agonists and active pacing (with frequencies just above spontaneous rate) as accelerators of oxygen consumption rate. The overall relationship between frequency of stimulation and oxygen consumption is a bit more complex, as revealed by computational modeling below.

### In silico analysis elucidates electro-mechano-energetic coupling in hiPSC-CMs

To address the goal of examining the utility of HT label-free long-term optical measurements of pericellular oxygen for assessing adrenergic drugs, we performed a complementary computational analysis. We modified a recent model of electro-mechano-energetic coupling in hiPSC-CMs [22] to provide a more accurate estimation of OCR, **Table 2**. In **Fig. 7A**, we compare electrical, mechanical and oxygen consumption responses under spontaneous (∼0.6 Hz) and paced (0.7 and 1 Hz) activity in hiPSC-CMs. Under spontaneous beating, the model shows that diastolic levels of OCR constitute 4.75% of peak oxygen consumption. In line with experimental data and other computational analysis, OCR is determined largely by myofilament activity, with additional contributions by non-phosphorylating respiration (proton leak + maintenance), sodium-potassium ATPase, SERCA pump and PMCA, detailed in **Fig. 7B**. Our simulations reveal a close relationship between oxygen consumption rate (OCR) and mechanical tension under the conditions studied, **Fig. 7A**. A slight increase in rate (from ∼0.6 to 0.7 Hz) leads to slight increase in peak tension and corresponding increase in OCR. Further increase in pacing rate (to 1 Hz) drops peak tension and peak OCR concurrently. This suggests a non-monotonic force-frequency relationship in the hiPSC-CMs, as seen also in cultured neonatal rat cells [44] and in engineered cardiac tissues [45]. This non-monotonic or largely negative force-frequency relationship driving OCR responses is further examined when modeling isoproterenol effects on hiPSC-CMs, **Fig. 8**.

**Figure 7.**
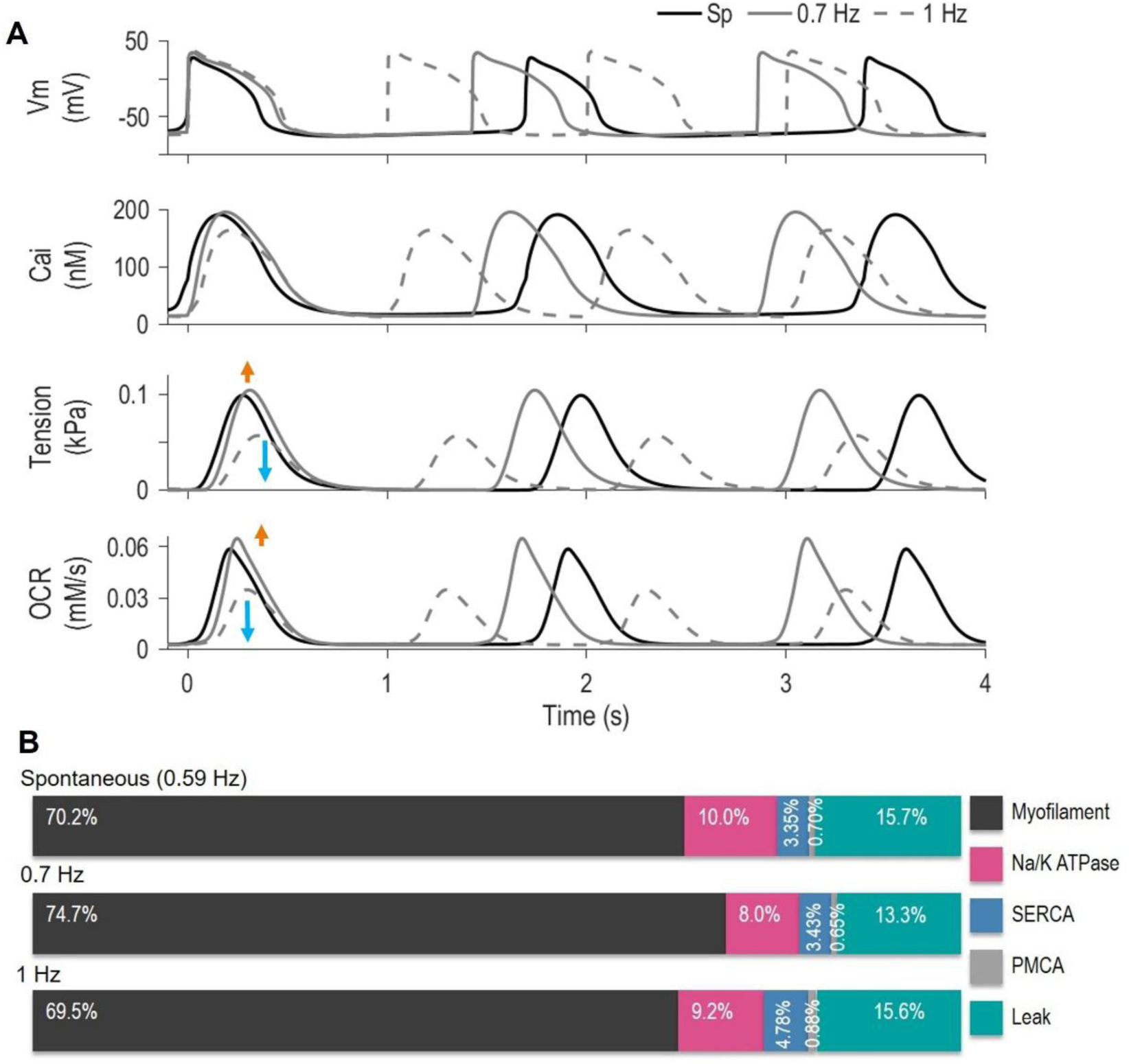
Computer simulations with an updated electro-mechano-energetic human iPSC-CM model derived from [22, 38]. **A.** Responses for membrane voltage, intracellular calcium, tension and OCR under spontaneous activity, 0.7 Hz pacing and 1 Hz pacing. Orange and blue arrows indicate the change (increase and decrease) with respect to spontaneous responses for 0.7 Hz and 1 Hz pacing, respectively. Based on the OCR traces in (A), we quantified the following OCR time integrals: 0.673 mM/min for spontaneous condition, 0.794 mM/min for 0.7-Hz pacing, and 0.678 mM/min for 1-Hz pacing. **B**. Summary of the relative contributions from myofilament ATPase, SERCA, sodium-potassium ATPase, PMCA, and non-electrophysiological ATP consumption (leak) to the total OCR for each pacing condition in panel A.

**Figure 8.**
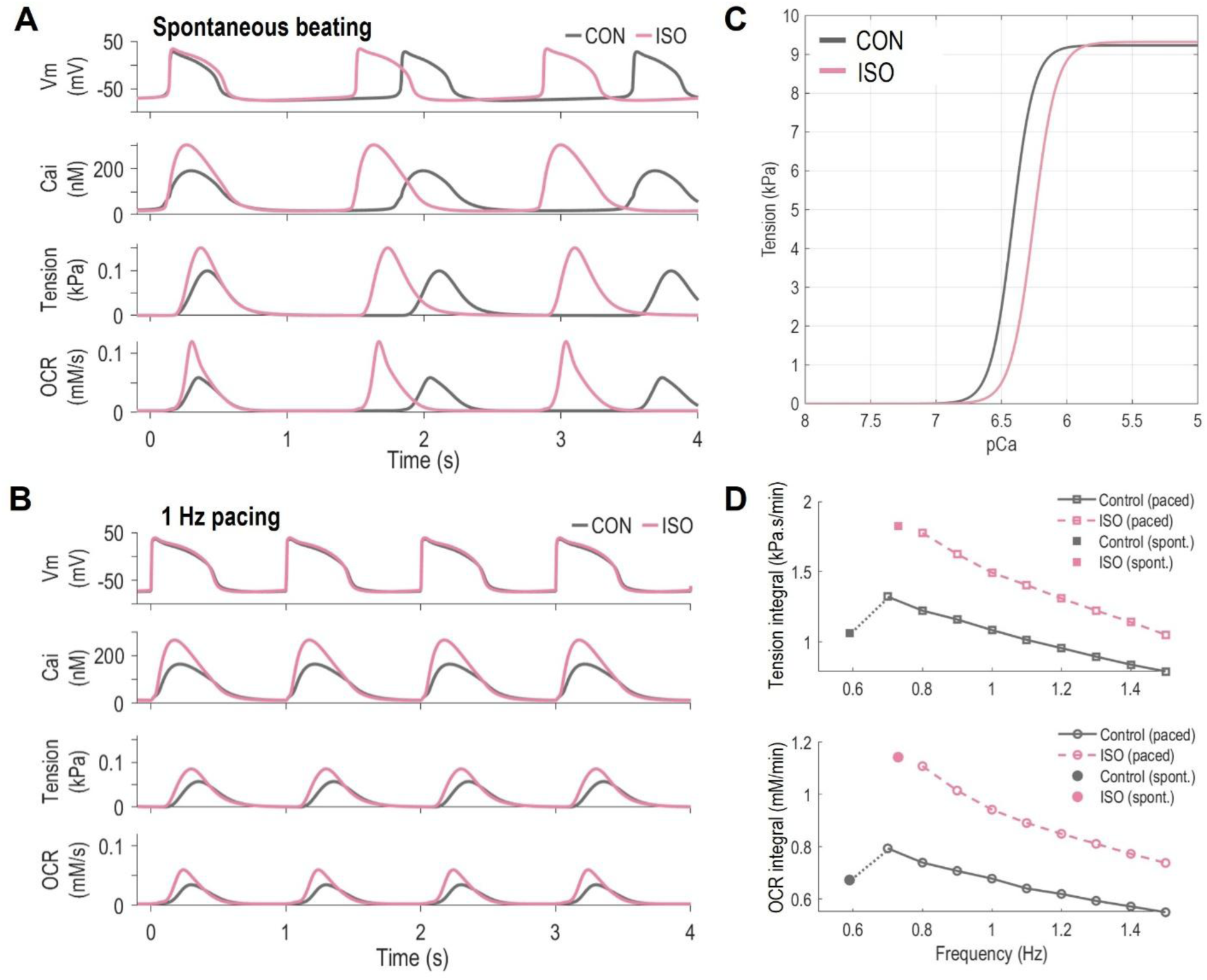
Computational modeling of responses to isoproterenol (ISO) in human iPSC-CMs showing chronotropic and inotropic effects with impact on tension development and OCR. A-B. Simulated membrane voltage, intracellular calcium, tension and OCR for control (CON) and ISO treatment under spontaneous activity and 1 Hz pacing, respectively. **C.** Calcium sensitivity of tension development for ISO vs. CON. **D**. Frequency dependence of time integrals of tension (top) and OCR (bottom) for ISO vs. CON. Left-most points (filled symbols) show values for spontaneous activity, the rest are paced.

### Computational modeling of hiPSC-CMs’ response to adrenergic agonists (ISO) reveals a key role of inotropy to increased OCR

To advance the mechanistic understanding of how adrenergic drugs modulate the coupled electrical, mechanical, and metabolic states of hiPSC-CMs, we simulated the effects of ISO administration. Functional effects induced by PKA-dependent phosphorylation on multiple targets were implemented as detailed in **Table 3A**. The resulting effects on hiPSC-CMs’ electrophysiology, mechanics, and energetics were validated against an independent set of experimental data describing changes in beat rate, action potential and calcium transient duration, and pericellular oxygen from this study as well as parameters from other studies [35–37], **Table 3B**.

Upon ISO administration, the model reproduced both the chronotropic (increase in rate) and the inotropic (increase in tension) effects induced by adrenergic stimulation, **Fig. 8A**. Increasing rate alone without corresponding inotropic effects of the drug could not explain the experimental OCR results. Our simulations show that the inotropic effects are still seen in rate-controlled (paced) conditions, **Fig. 8B**. A key aspect of ISO-induced effects is the increased calcium sensitivity (higher pCa) of tension development, **Fig. 8C**. ISO-induced changes in mechanical tension (increase) are closely followed by changes in OCR (increase) over a range of frequencies, **Fig. 8D**. Based on these non-monotonic, primarily negative force-frequency relationships (and the corresponding OCR-frequency curves) in the studied hiPSC-CMs, we surmise that the ISO effects on OCR in hiPSC-CMs (seen in **Figs. 3, 5, 6**) are likely largely due to ISO’s inotropic action (increase in tension compared to control). During spontaneous activity, this OCR increase is likely aided by the smaller chronotropic effect of ISO, **Figs. 4** and **6**.

### Experiments and computational modeling of hiPSC-CMs’ response to excitation-contraction uncouplers (BLEBB) underscore the dominant contribution of contraction to OCR

Focusing on the close links between tension and OCR, we decided to investigate the extreme case of excitation-contraction uncoupling induced by BLEBB administration, **Fig. 9**. First, we conducted a set of *in vitro* experiments to quantify BLEBB effects on OCR in hiPSC-CMs, **Fig. 9**. In this static culture, we found that BLEBB produced a long-lasting response characterized by decreased depletion of pericellular oxygen and reduced OCR, evident from about 0.5 h to 20 h post-administration. We then simulated BLEBB effects by implementing the computational methodology previously described in [38]. The parameter modifications summarized in **Table 4A** enabled the reproduction of key experimental output metrics (**Table 4B**), including decreased myofilament calcium sensitivity (pCa) and tension-calcium cooperativity (nHill) consistent with [46]. Our simulations reproduced also the significant reduction in peak tension observed experimentally in [35] and in OCR (this study), **Table 4B**.

**Figure 9.**
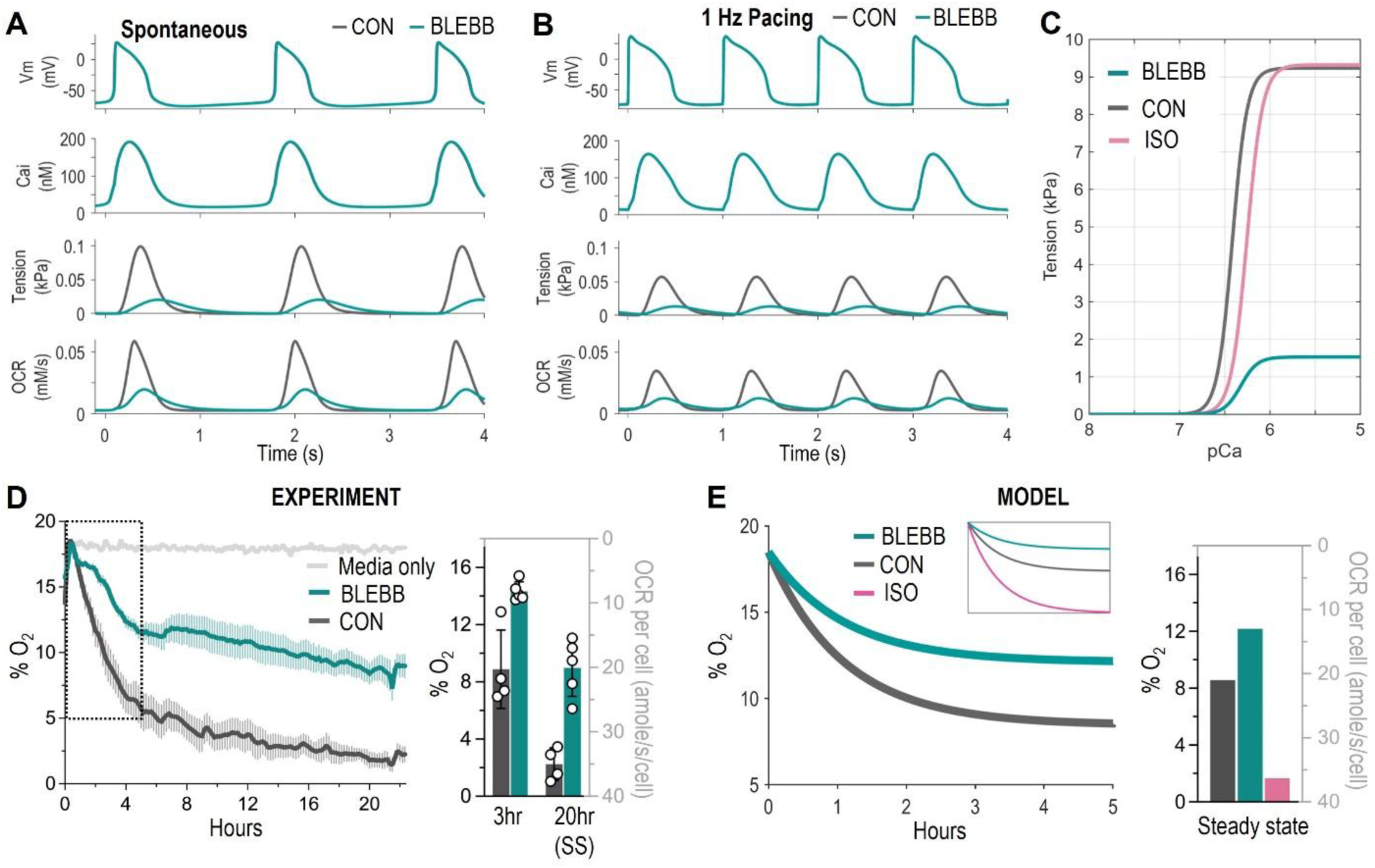
Computational modeling and experimental results for responses to Blebbistatin (BLEBB) in human iPSC-CMs with impact on tension development and OCR. A-B. Simulated membrane voltage, intracellular calcium, tension and OCR for control (CON) and BLEBB treatment under spontaneous activity and 1 Hz pacing, respectively. **C.** Calcium sensitivity of tension development for BLEBB, ISO and CON (note the shift in pCa and drop in maximum tension developed for BLEBB). **D**. Experimental results for pericellular oxygen over 22 hours for BLEBB (2.5mM) vs. CON with media only control, n = 5, data shown are averaged curves with SEM. Bar plots show the individual sample measurements for 3 h and 22 h (steady state) after BLEBB application. The y-axis on the left shows pericellular oxygen levels (%) and on the right estimated OCR per cell (amole/s/cell). **E.** Simulation results for the experiment from panel D (see boxed area) with BLEBB treatment, inset shows results including CON, BLEBB and ISO. Bar plots on the right display the steady state values from the curves for CON, BLEBB and ISO (in agreement with Fig. 3B-D and Fig. 9D).

Compared to control, BLEBB application led to a drop in maximum tension of about 80% in both spontaneous and paced conditions, **Fig. 9A-C**. Consistent with our *in vitro* results, BLEBB simulations also yielded a significant reduction in OCR. Notably, the model qualitatively reproduced our experimental measurements of pericellular oxygen depletion dynamics and steady-state OCR values over a 5-h interval, **Fig. 9D-E**. Together, these findings suggest that the experimental effects seen with adrenergic antagonists (**Figs. 3-6**) likely reflect a combination of reduced contractile demand (as in BLEBB experiments) and decreased beat rate, especially for propranolol that often silenced activity.

## DISCUSSION

In this study we explored the question if label-free chronic optical observation of oxygen consumption in cardiomyocytes can be used to successfully differentiate the action of rate- and contractility-modulating drugs (adrenergic agonists and antagonists). Such approach has multiple advantages, including: 1) simplicity (can be implemented in a very low cost, compact format), 2) scalability (all measurements reported here are in standard 96-well plates with a whole-plate readout), 3) no interference with cellular function (no dyes or modifications are needed), 4) compatibility with long-term monitoring (suitable for days to months of cellular observation as various treatments are applied), and 5) sophistication of captured parameters (multiple metrics can be extracted that quantify metabolic function from the relatively simple measurements).

In previous work, we have validated that the method can be used to report true absolute values of oxygen, just as a classic electrochemical Clark electrode can, with the benefit that it can be calibrated to track changes in oxygen in the tight pericellular space (immediately surrounding the cells), where Clark electrodes are not applicable [13, 14]. We have shown the method to be able to detect quantitative differences in oxygen consumption in human cardiac fibroblasts and human iPSC-cardiomyocytes and distinguish states of normoxia, hyperoxia and hypoxia[14]. Furthermore, a substantial increase in metabolism could be tracked in iPSC-CMs over five days in culture, with clear differences depending on the conditions of growth, with the cells grown in conditions close to normoxia showing the greatest improvement [14]. Our results of cardiac cell oxygenation in HT plates brought about important insights and some surprising observations, e.g. we found hyperoxic state for cardiac fibroblast growth and for sparse cardiomyocyte cultures, yet highly hypoxic conditions were observed for static cardiomyocyte cultures when grown in dense syncytia. These previous investigations gave us confidence that this HT approach has great potential in tracking metabolic cardiac cell responses over time and can be leveraged in drug development.

With the knowledge that cardiomyocytes are highly metabolically active cells with their electromechanical activities using up to 80% of the oxygen, we focused on rate-and contractility-modulating drugs and their signature in the measured oxygen consumption curves. A cross-validation of the activity of the adrenergic agonists and antagonists on the iPSC-CMs was done in the same samples (from which optical oxygen measurements were obtained) using HT all-optical cardiac electrophysiology. Future investigations can provide more time-resolved direct validation using concurrent measurements of oxygen and electrophysiology over time, as the sequential measurements presented here bear limitations due to the dynamic nature of drug-induced changes. However, even for the single time-point measures of rate, our approach here yielded convincing correlation between electromechanical activity and metrics extracted from the oxygen consumption curves. Using optogenetic pacing over several hours, we confirmed heart rate can affect the level of depletion of pericellular oxygen in the iPSC-CMs. In terms of useful metrics for future drug testing studies, we found that adrenergic antagonists, such as β-blockers, are best distinguished by the steady-state levels of pericellular oxygen within several hours of treatment (and depending on dose, perhaps earlier). Adrenergic agonists were best identified by the time to reach 5% (or hypoxic levels of 2%) of pericellular oxygen, which directly relates to the initial rate of depletion. The same was true for differentiating the actively paced cells from spontaneously active cells.

Given the complexity of linking adrenergic modulation to changes in electrophysiology, contractility, and oxygen consumption, we complemented our *in vitro* approach with computational modeling. Extending recent work [22–24, 38], we updated the model of electro-mechano-energetic coupling in hiPSC-CMs by improving OCR estimation and introducing a novel parametrization of the effects of PKA signaling. To our knowledge, the latter represents the first implementation of this kind in hiPSC-CM models. Our *in silico* analysis helped obtain a better understanding of the multifaceted effects of adrenergic agonists, specifically ISO, on hiPSC-CMs. The simulations revealed that the inotropic action of ISO is a stronger contributor to the OCR increase and likely dominates its chronotropic effects. Furthermore, using the excitation-contraction uncoupler BLEBB in both experiments and simulations, we illustrated the tight coupling between mechanical tension and OCR. As expected, BLEBB caused a marked reduction in peak tension under both spontaneous and paced conditions, accompanied by a substantial decrease in peak OCR. Our model also captured the experimentally observed dynamics of pericellular oxygen depletion following suppression of contractility in a semi-quantitative manner.

To our knowledge, no prior studies have attempted to simulate such a comprehensive response in hiPSC-CMs, particularly by integrating action potentials, calcium cycling, force generation, and oxygen consumption within a unified framework. However, this breadth comes with limitations. Specifically, our framework only explicitly includes four ATP-consuming processes (i.e., myofilament, SERCA, sodium-potassium ATPase, and PMCA), while additional ATP sinks contributing to basal metabolic demand are not modeled individually. Non-electrophysiological ATP consumption (e.g., biosynthesis, protein turnover, mitochondrial maintenance) is subsumed into the OCR_leak_ term that is treated as constant despite its known dependence on mitochondrial membrane potential, substrate availability, and maturational state [30, 31, 47]. While the commercial cell line iCell2 used and modeled here as well as the proprietary culture medium have been optimized metabolically [48], further maturation efforts are actively ongoing. Lastly, our framework does not include an explicit representation of ATP production pathways (e.g., glycolysis, fatty acid oxidation, oxidative phosphorylation dynamics). While these simplifications make the model particularly well suited to connect electromechanical activity to OCR under controlled baseline conditions, future extensions will be needed to capture metabolic heterogeneity and stress-response behaviors more fully. Nevertheless, validation of this *in silico* electro-mechano-energetic representation using published datasets together with our experimental results represents a significant step toward a comprehensive, multiparametric hiPSC-CM “digital twin”. This will enable mechanistic, quantitative interpretation of drug responses and support a broad range of future investigations, including informed microfluidics-based control of oxygenation for human-based platforms [49].

The effects of adrenergic modulators and other rate-, contractility- and metabolism-altering drugs are complex and may involve spatial effects, i.e. effects on cell-cell coupling and on conduction velocity (dromotropic action). Such spatially-distributed effects are important arrhythmia determinants and are not captured in the experimental nor computational analysis presented here. Since scalability was our focus, we used only space-averaged measures per sample and a single-cell computational model to achieve that. Both the all-optical experimental approaches and the modeling are well suited for future spatially-resolved investigations to quantify such drug effects in multicellular function, with some sacrifice of scalability. Future work can also expand the panel of metabolic compounds tested and use the entire shape (temporal profiles) of pericellular oxygen curves in conjunction with machine learning algorithms [50] for building classifiers and more sensitive predictors of drug action in cardiomyocytes, as well as for quality control of various metabolic maturation strategies [51, 52] for human iPSC-CMs.

In conclusion, with high interest in development of rate- and contractility-control drugs and metabolic modulators, we envision the reported hybrid high-throughput experimental and computational methodology to be widely applicable, with high translational potential, especially with the focus on human-based preclinical investigations [11].

## Author contributions

EE and WL designed the experimental study. WL conducted most experiments, analyzed and plotted the data and co-wrote the first draft. MAF and SM worked on the computational modeling. MAF formulated and implemented all computational models, run simulations, analyzed the results and prepared a writeup. DM helped with calibration, setup and validation of the optical oxygen system as well as the methodology for cutting and mounting of optical oxygen sensors. MRP conducted some of the experiments for the revision, analyzed data, generated plots and aided in writing. YWH developed the all-optical electrophysiology systems, the optogenetic stimulation setup and aided in data analysis. MWK and ZL helped with supervision and research resources for the validation of the optical oxygen measurements, optimization of parameters and experimental design. SM provided input for the computational modeling, supervision and research support. EE supervised all aspects of the project, secured funding and co-wrote the initial draft. All authors provided input on the manuscript and its revision.

## Data Availability

The experimental data underlying this study are available in the published article. The Matlab model code is available for download at Zenodo (*link and DOI to be included upon publication*) and https://github.com/aminforouzandehmehr.

## Acknowledgements

This work was supported in part by grants from the NIH-NHLBI R01HL144157 (to EE and MWK) and from the NSF EFMA1830941 (to EE and ZL). SM was supported by grants from the NIH-NHLBI R01HL171057, R01HL171586, R01HL176651. MAF was also supported by Finnish Cultural Foundation (SKR) under grant number: 00241356. Biorender was used to create parts of figures 1, 2, 3 and 6.

